# HIV-1 Reverse Transcriptase interactions with Long-acting NNRTI, Depulfavirine (VM1500A)

**DOI:** 10.64898/2026.04.06.715899

**Authors:** A.A. Snyder, I.L. Kaufman, C.J. Risener, K.A. Kirby, S.G. Sarafianos

## Abstract

Non-nucleoside reverse transcriptase inhibitors (NNRTIs) are key components of combination antiretroviral therapy (ART) for the treatment of human immunodeficiency virus type 1 (HIV-1) infection, binding an allosteric pocket of reverse transcriptase (RT) and inhibiting viral replication. Although second-generation NNRTIs have improved potency and resistance profiles compared to first-generation NNRTIs, the continued emergence of resistant viral strains and the need for long-acting therapeutic options underscore the importance of developing next-generation compounds. Depulfavirine (VM1500A) is a potent NNRTI being developed as a long-acting formulation. Its prodrug, elsulfavirine (ESV), is approved for HIV-1 treatment in Eurasian countries as a once-daily oral regimen and has demonstrated favorable antiviral efficacy, pharmacokinetics, and tolerability in clinical studies. Here, we report the 2.4 Å crystal structure of HIV-1 RT in complex with depulfavirine, revealing an extended binding conformation within the NNRTI pocket that reaches from the back of the binding pocket to the entrance. These interactions may shed light on mechanisms of resistance to the F227C mutation, with and without V106 substitution, and Y188L. Notably, depulfavirine maintains potent inhibition of common NNRTI-resistant RT variants, including K103N and Y181C. Combination studies of ESV with antivirals from diverse inhibitor categories demonstrated additive or near-synergistic activity with islatravir (ISL), cabotegravir (CAB), lenacapavir (LEN), and tenofovir (TDF). These findings highlight the broad resistance profile and potential of the depulfavirine combination.

## Introduction

As of 2024, an estimated 40.8 million people are living with human immunodeficiency virus type 1 (HIV-1) worldwide (1). While there is currently no cure for HIV, antiretroviral therapies (ART) have significantly improved disease management by reducing viral load to undetectable levels and improving patient outcomes. Despite this success, several limitations remain, including challenges with patient adherence to strict dosing regimens, treatment-related side effects, high costs, and reduced efficacy due to the emergence of drug resistance. As a result, treatment-experienced i(2)ndividuals often require additional therapeutic options. Therefore, there remains a need to develop new antivirals, particularly those with long-acting potential. The standard of care for HIV treatment is a once-daily oral pill, but novel long-acting treatment options are being developed (2). Currently, Cabenuva (cabotegravir (CAB) + rilpivirine (RPV)) and Sunlenca (lenacapavir (LEN)) are the only FDA-approved long-acting antiviral options for HIV treatment, while Yuztugo (LEN) and Apretude (CAB) are approved for pre-exposure prophylaxis (PrEP) (3). These treatments are highly successful and represent major advances in long-acting therapy. Still, there remain far fewer long-acting options than daily oral options; additionally, these treatments have only been available since 2021 (2).

The first U.S. Food and Drug Administration (FDA) approved non-nucleoside reverse transcriptase inhibitor (NNRTI) was nevirapine (NVP) in 1996, followed by delavirdine in 1997 and efavirenz (EFV) in 1998 (3). First-generation NNRTIs marked a breakthrough in HIV treatment; however, their clinical usefulness was soon challenged by the emergence of drug-resistant viral strains (4). In response, second-generation NNRTIs such as etravirine (ETR) were developed and approved in 2008, offering improved flexibility to accommodate resistance mutations and reduce adverse side effects (5, 6). NNRTIs target an allosteric site at the base of the thumb of HIV-1 reverse transcriptase (RT) enzyme to inhibit viral DNA polymerization or extension (7, 8). The NNRTI binding pocket is formed by the key residues: L100, K101, K103, V106, T107, V108, V179, Y181, Y188, V189, G190, F227, W229, L234, and Y318 of p66 and E138 of p51 and is only formed in the presence of an NNRTI (9). Currently, five NNRTIs are approved for use in the United States: NVP, EFV, ETR, doravirine (DOR), and RPV, the latter of which is also formulated for long-acting administration (3). Despite their success, there is still a need to develop new NNRTIs with improved resistance profiles and additional options suitable for long-acting delivery (2, 10).

One promising candidate is elsulfavirine (ESV), the prodrug of depulfavirine (VM1500A), 2-(4-bromo-3-(3-chloro-5-cyanophenoxy)-2-fluorophenyl)-N-(2-chloro-4-sulfamoylphenyl)acetamide, developed by Viriom (Scheme 1) (11). Elsulfavirine has been approved in Russia and the Eurasian Economic Community (EurAsEC) countries under the name Elpida since 2017, while depulfavirine is currently in clinical trials as a long-acting injectable (e.g., NCT05204394). *In vitro* studies have demonstrated that ESV has potent activity against wild-type HIV-1, with an IC_50_ of ∼ 1 nM, and maintains efficacy (IC_50_ < 100 nM) against over 90% of tested NNRTI-resistant clinical isolates, including those with common resistance mutations such as K103N and Y181C. Viral passaging experiments with increasing ESV levels revealed that multiple mutations, such as V106I/A combined with F227C or Y188L, are needed to confer significant resistance. Notably, these multi-mutation pathways often result in decreased viral fitness compared to wild-type strains, suggesting that ESV and depulfavirine may present a higher genetic barrier to resistance than current NNRTIs (11). Additionally, the t_1/2_ of depulfavirine is approximately 5.4 to 7.4 days, depending on the dose, which is substantially longer than that of most NNRTIs in clinical use (12). These results highlight the potential of ESV for long-acting use (13).

**Scheme 1.**
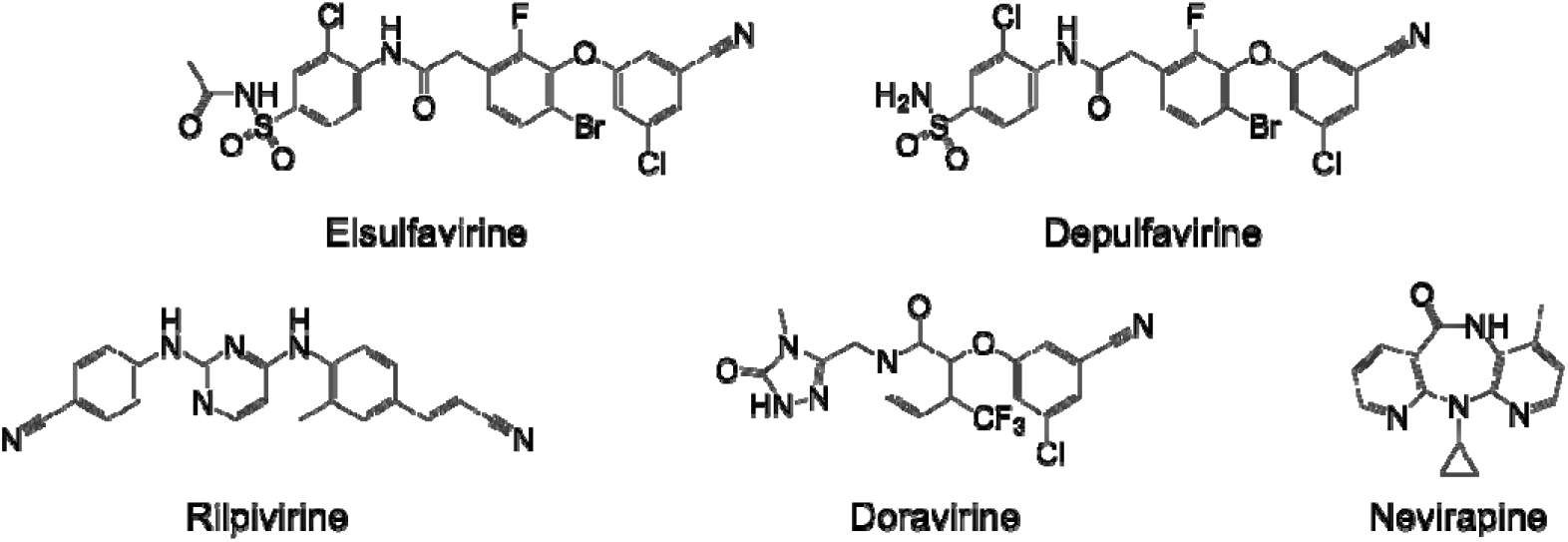
Chemical Structures of NNRTIs: Elsulfavirine (ESV), Depulfavirine (VM1500A), Doravirine (DOR), Rilpivirine (RPV), and Nevirapine (NVP). ESV is the prodrug to the active form of depulfavirine.

Currently, the structural basis of depulfavirine binding, resistance tolerance, and combination profile has not been fully explored. In this study, we determined the 2.4 Å X-ray crystal structure of HIV-1 RT in complex with depulfavirine, revealing the key molecular interactions that underlie its inhibitory mechanism and resistance profile. Structural analysis shows that depulfavirine adopts a binding mode distinct from that of both first-generation NNRTIs, such as NVP, and second-generation inhibitors, including RPV. Its binding most closely resembles that of the clinically relevant NNRTI, DOR, although several structural differences may contribute to the observed variations in resistance profiles. Defining these molecular determinants provides insight into how depulfavirine accommodates resistance-associated changes within the NNRTI-binding pocket and informs the design of next-generation long-acting NNRTIs. We further evaluated the impact of clinically relevant resistance mutations *in vitro* and assessed combination activity with potential long-acting antiretroviral partners, observing additive interactions between elsulfavirine (ESV) and multiple agents.

## Methods

### Protein Expression and Purification

#### Crystallography construct

p66 and p51 RT genes from BH10 HIV-1 were cloned into the pETDuet-1 vector (Novagen) using restriction sites NcoI and SacI for the p51 subunit and SacII and AvrII for the p66 subunit. A hexahistidine tag and a 3C protease recognition sequence were added to the N-terminus of the p51 subunit for purification purposes. The vectors were then transfected into BL21 cells, and the RT genes were expressed. RT was then purified using nickel affinity chromatography and MonoQ anion-exchange chromatography (14, 15).

#### Biochemistry construct

HIV-1 RT (p6HRT-prot) was expressed in JM109 cells (Agilent, Catalog: 200235) and purified as previously described by nickel affinity chromatography (14).

### Crystallization of the NNRTI/RT complex

RT crystals were formed by hanging drop trays at 4 °C in solutions containing 11-12% PEG 8,000, 100 mM ammonium sulfate, 0.05 M bis-tris propane, pH 6.4, 5% ethylene glycol, and a 2-fold molar excess of VM-1500A (synthesized by Viriom Inc.). Crystals were cryoprotected with 22% ethylene glycol and 20% PEG 8000 for up to 5 minutes and then flash-cooled in liquid nitrogen.

### Data Collection and Determination

Data were collected on a DECTRIS PILATUS3 6M detector at the Advanced Photon Source (APS) beamline 23-ID-D, Argonne National Laboratories (<u>Table S2</u>). Datasets were processed using XDS (16) and indexed in C121 (*a* = 163.05 Å, *b* = 73.87 Å, and *c* = 108.64 Å) space group with one RT/NNRTI complex per asymmetric unit. The crystal structure was solved with molecular replacement using ID 4G1Q from the Protein Data Bank (PDB) and PHASER (17) within the CCP4 package (18). Several rounds of manual rebuilding using Coot (19) and refinement with the PHENIX (20) and CCP4 (18) were performed.

### Cell Culture

DMEM/High Glucose (Cytiva HyClone^™^ Dulbecco’s Modified Eagle’s Medium) with 10% FBS, and 50 U/mL penicillin/50µg/mL streptomycin (Thermo Fisher) was used to culture HEK293T/17 (21) and TZM-green fluorescent protein (GFP) (22) (received from Massimo Pizzato) cell lines for subsequent experiments.

Single-nucleotide mutagenesis was used to mutate F227C in the RT protein of NL4-3ΔenveGFP plasmids using NEB HiFi Assembly (Biolabs). DNA was sequenced to verify the presence of each mutation. Then, VSVG-pseudotyped virus stocks were prepared. 5 μg of pNL4-3Δenv eGFP mutant and WT plasmids (from Bei Resources ARP-11100) were each transfected with 1.5 μg of pVSVG (Bei Resources ARP-4693) in HEK293T cells using Xtreme-GENE HP transfection reagent (Roche). 48 h post-transfection, the supernatant was collected, centrifuged for 5 min at 2,000 rcf, and filtered through a 0.22 μm polyethersulfone (PES) membrane (Millipore) to remove residual cells.

### Synergy: ESV+ LEN/CAB/TDF (WT)

TZM-GFP cells and 2-fold serially dilutions of ESV (Viriom) with ISL (Life Chemicals), LEN (Medchem Express), CAB (Selleck Chemicals), or TFV (Bei Resources) in a matrix format were plated together in 96-well plates and incubated for 24 h. After 24 h, the cells were infected with a mixture of DEAE-dextran (1 μg/mL final concentration) and NL4-3ΔenveGFP WT VSVG pseudotyped virus. After infection, plates were incubated for an additional 48 h. Cytation 5 (BioTek) was used to image GFP-positive cells, which were counted using Gen5 v3.15 (BioTek). To determine percent inhibition, two technical replicates were averaged, and dose-response data were normalized to no-drug controls. Experiments were conducted in triplicate and analyzed using SynergyFinder Plus software (23).

### Polymerase inhibition assays

The DNA template was annealed to a 5′-Cy3–labeled DNA primer at a 3:1 molar ratio (Td31: 5′-CCA TAG CTAG CAT TGG TGC TCG AAC AGT GAC; Pd18-P0: 5′-Cy3 GTC ACT GTT CGA GCA CCA) (Integrated DNA Technologies). For primer extension assays, the resulting DNA/DNA hybrid (20 nM) was incubated at 37 °C with either wild-type (WT) or mutant HIV-1 reverse transcriptase (RT) (Y181C, V106A, F227C, V106A/F227C, K103N, or E358K) at a final concentration of 20 nM in RT buffer containing 50 mM Tris (pH 7.8) and 50 mM NaCl. Reactions were initiated by adding 6 mM MgCl_2_ after introducing varying concentrations of depulfavirine, and the final reaction volume was 20 μL. All dNTPs were present at a final concentration of 1 μM. After 15 min, reactions were quenched by adding an equal volume of 100% formamide containing trace amounts of bromophenol blue. Reaction products were separated on 15% polyacrylamide gels containing 7 M urea and visualized using a Typhoon FLA 9000 phosphorimager (GE Healthcare, NJ). Bands corresponding to fully extended products were quantified using AzureSpot Pro. Data from at least three independent experiments were analyzed and plotted as the percentage of full extension (extended product/total product). IC_50_ values for depulfavirine were determined by nonlinear regression using a log(inhibitor) versus response variable-slope (four-parameter) model in GraphPad Prism 10 and are reported as mean ± standard deviation.

## Results/Discussion

### Structure

The structure of HIV-1 RT bound to depulfavirine provides insights into the molecular basis of this NNRTI inhibition of viral synthesis (Figure 1). All NNRTIs bind to a hydrophobic pocket at the base of the thumb, adjacent to the polymerase active site. The binding pose and interactions observed in this study differ in several key interactions compared to other HIV-1 NNRTIs, providing insights into the high potency and resistance profile of this molecule.

**Figure 1:**
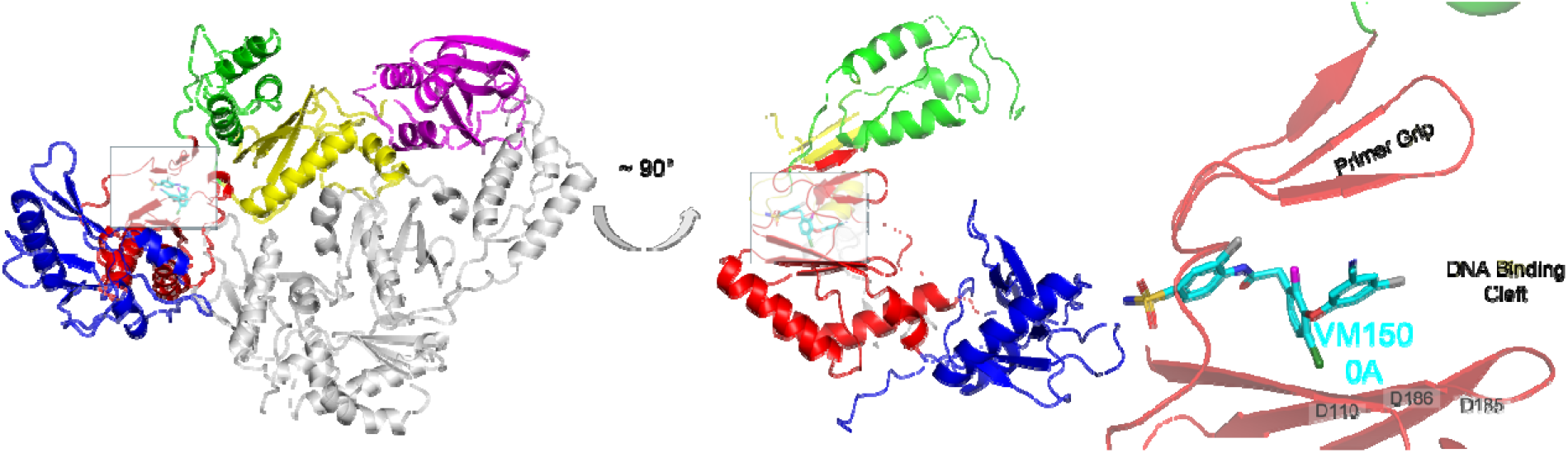
X-ray crystal structure of the HIV-1 RT: depulfavirine complex. The p51 subunit is shown as white and the p66 subunit as multicolor: fingers (blue), palm (red), thumb (green), connection (yellow) subdomains, and RNase H domain (purple). Depulfavirine (VM1500A) shown in cyan and colored by atom: Cl in gray, F in magenta, Br in green, S in yellow, O in red, and N in blue. This structure has been deposited in the PDB under accession code 7TAZ.

Depulfavirine makes key interactions with residues K101, K102, K103, K104, S105, V106, V108, Y188, P225, F227, W229, L234, H235, P236, and Y318 (Figure 2). At the entrance of the pocket, the sulfonamide group forms contacts with the two prolines, P225 and P236, as well as with the backbone atoms of the lysine-rich region (K101–K104). Within this cluster, three distinct hydrogen bonds are made with K102, K103, and K104, anchoring the inhibitor at the mouth of the NNRTI-binding pocket. Depulfavirine engages in multiple hydrophobic and van der Waals interactions with V106, V108, F227, W229, L234, and Y318, thereby positioning the ligand in an extended conformation. Depulfavirine extends to the back of the pocket, interfacing with W229, and makes key hydrophobic interactions with V106, V108, F227, L234, and Y318. Importantly, depulfavirine also forms a π–π interaction with Y188, a hallmark contact observed in several highly potent NNRTIs. Together, these interactions define the binding mode of VM1500A.

**Figure 2:**
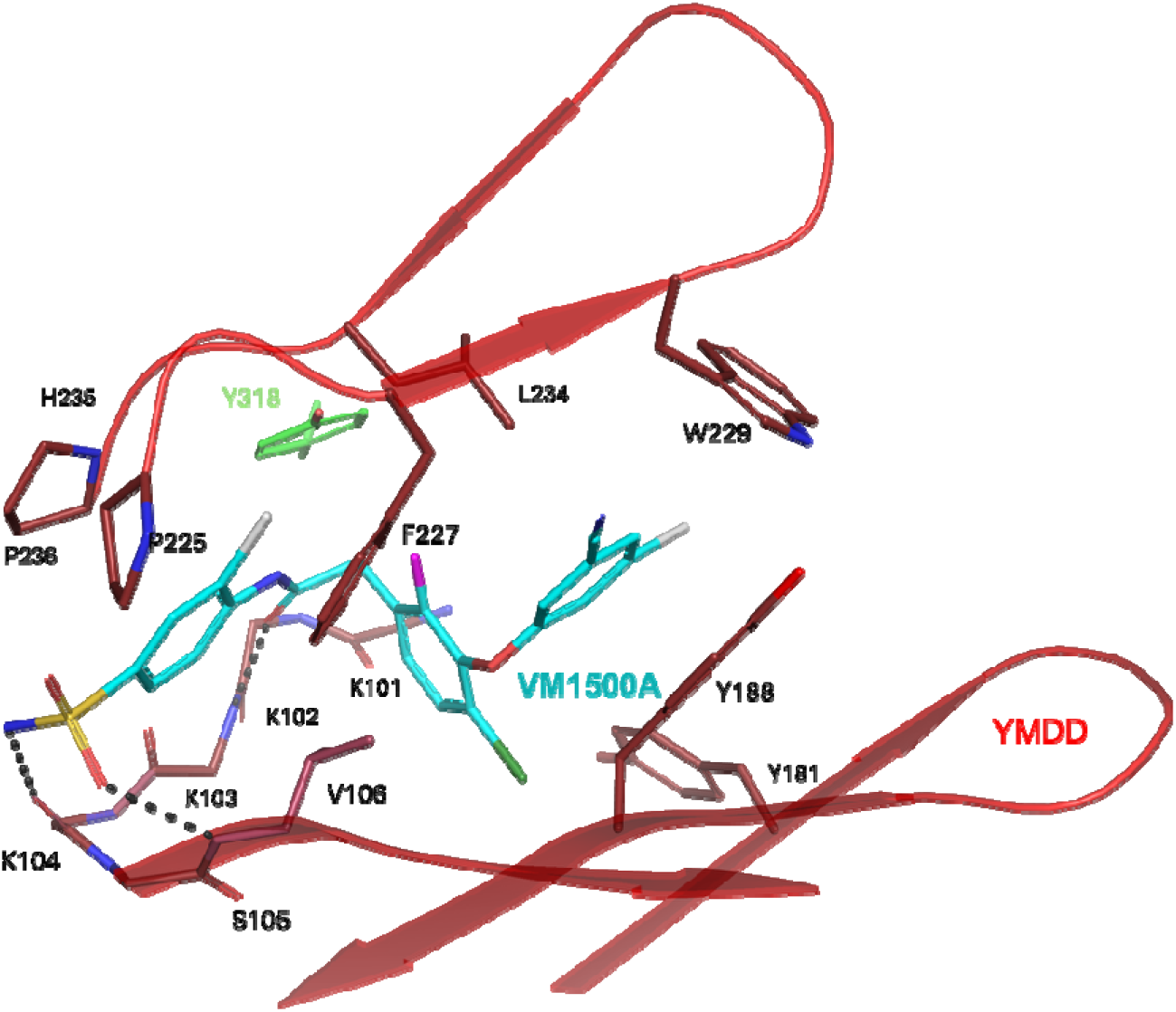
Depulfavirine binds at the NNRTI binding site of HIV-1 RT. Depulfavirine (VM1500A) is shown in cyan, and RT p66 is shown in red (palm) and green (thumb). Hydrogen bonds are shown in black dashes.

Like other second-generation NNRTIs, depulfavirine possesses a flexible three-ring system that enables greater molecular adaptability and improves tolerance to resistance-associated mutations compared with first-generation NNRTIs such as NVP (7). Here, we compare the binding interactions of depulfavirine with those of DOR, RPV, and NVP by aligning the crystal structures of the four inhibitors using the β-sheets surrounding the YMDD motif (Figure 3). Each NNRTI binds RT slightly differently and has unique interactions (Figure 3E, Sup Figure 4). Depulfavirine and DOR exhibit similar binding modes, extending into the deeper regions of the NNIBP (24); however, depulfavirine forms additional interactions near the pocket entrance (K101–S105) (Figure 3E). The cyanochlorophenol groups in both inhibitors form direct π–π interactions with Y188. Interestingly, Y181, a residue frequently associated with resistance, rotates away from Y188 and does not directly interact with either depulfavirine or DOR. Although Y181 adopts the same general orientation in both complexes, the aromatic ring is slightly twisted in the depulfavirine structure, which may also impact E138 (Figure 3B).

**Figure 3.**
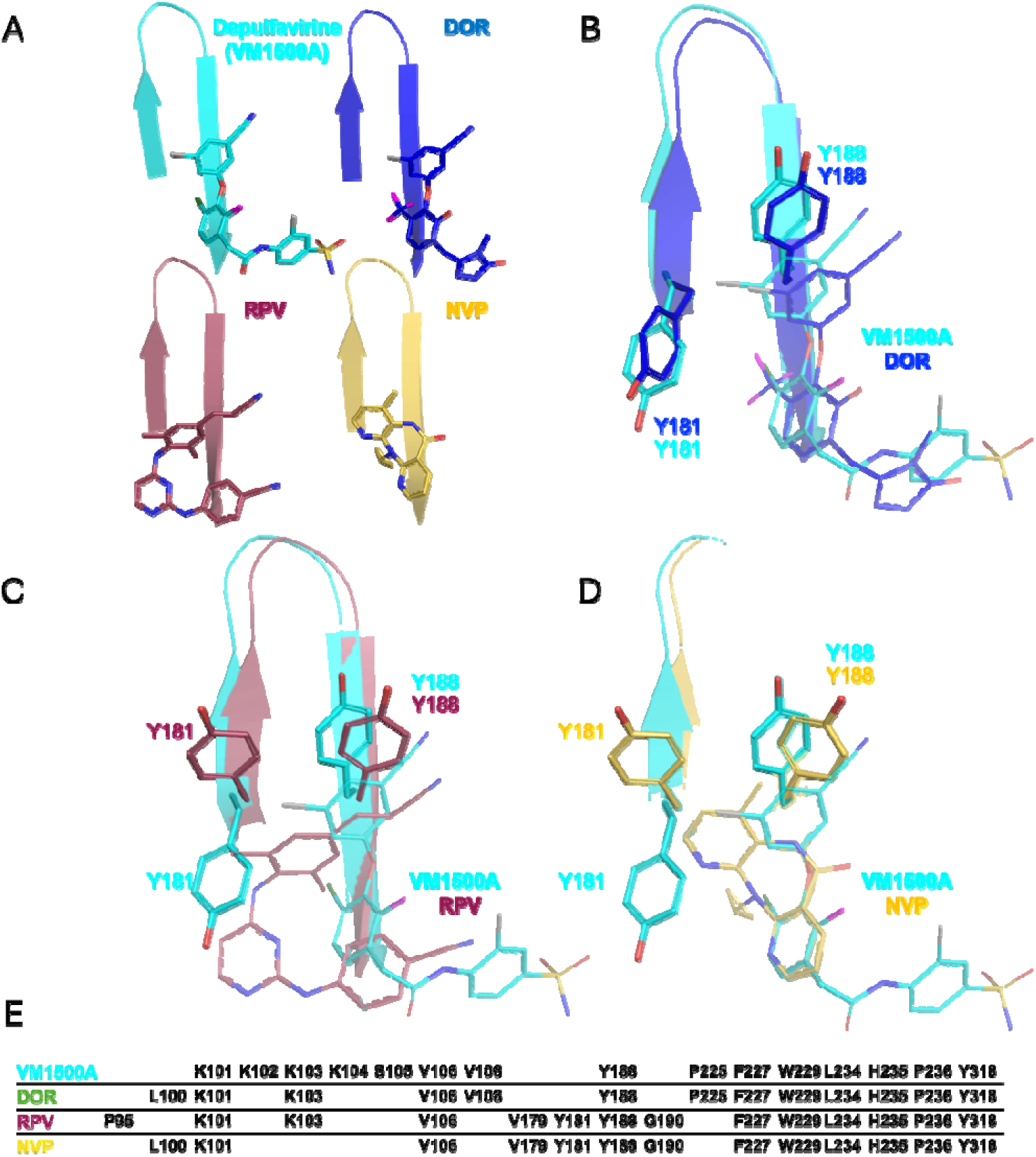
Distinct Conformations of Y181 Across NNRTI–RT Complexes Structural comparisons showing how different NNRTIs position Y181 within the HIV-1 reverse transcriptase (RT) NNRTI-binding pocket. (A) Overlay of depulfavirine (VM1500A) (cyan, PDB 7TAZ), DOR (blue, PDB 4NCG), RPV (rust, PDB 3MEE), and NPV (yellow, PDB 1VRT) aligned on the NNRTI-binding pocket. (B) Comparison of depulfavirine (VM1500A) and DOR interactions with RT. (C) Structural comparison of depulfavirine (VM1500A) and RPV. (D) Structural comparison of depulfavirine (VM1500A) and NPV. (E) Summary of key NNRTI–RT interactions within 3.7 angstroms.

Although RPV is also a second-generation NNRTI, it adopts a U-shaped conformation within the pocket, allowing simultaneous interactions with Y181 and Y188, aligning these residues in the same orientation (Figure 3C). (25) Similarly, NVP interacts with the same conformations of Y181 and Y188 (Figure 3D) (26). Notably, these shifts in Y181 positioning affect the local environment around E138 of p51, located at the p66/p51 interface of HIV-1 reverse transcriptase (27). Mutations at this site, particularly E138K, are associated with resistance to rilpivirine (RPV) and can influence RT structure, viral replication capacity, and enzyme stability (28, 29). NNRTIs, including RPV, have also been shown to promote RT dimerization within the Gag-Pol polyprotein and premature protease activation (30, 31).

### Resistance

Across a panel of clinically relevant NNRTI resistance mutations, ESV, the prodrug of depulfavirine, displays a distinct resistance profile compared with other approved NNRTIs (Figure 4) (4, 11, 24, 32). Most single mutations confer little to no reduction in susceptibility to ESV/depulfavirine, including L100I, K103N, V106A, Y181C, Y181I, G190A, and M230L, which remain within approximately two-fold of wild-type susceptibility in viral infectivity assays of RO-0335 (depulfavirine/VM1500A) (11). A modest reduction in susceptibility is observed with E138K (∼2-fold) (11). High-level resistance to depulfavirine is primarily associated with Y188L and mutations involving F227C (Supplemental Figure 1), particularly the V106A/F227C pathway and higher-order variants such as V106A/F227C/M230L, which exhibit >100-fold reductions in susceptibility (Figure 4) (11).

**Figure 4.**
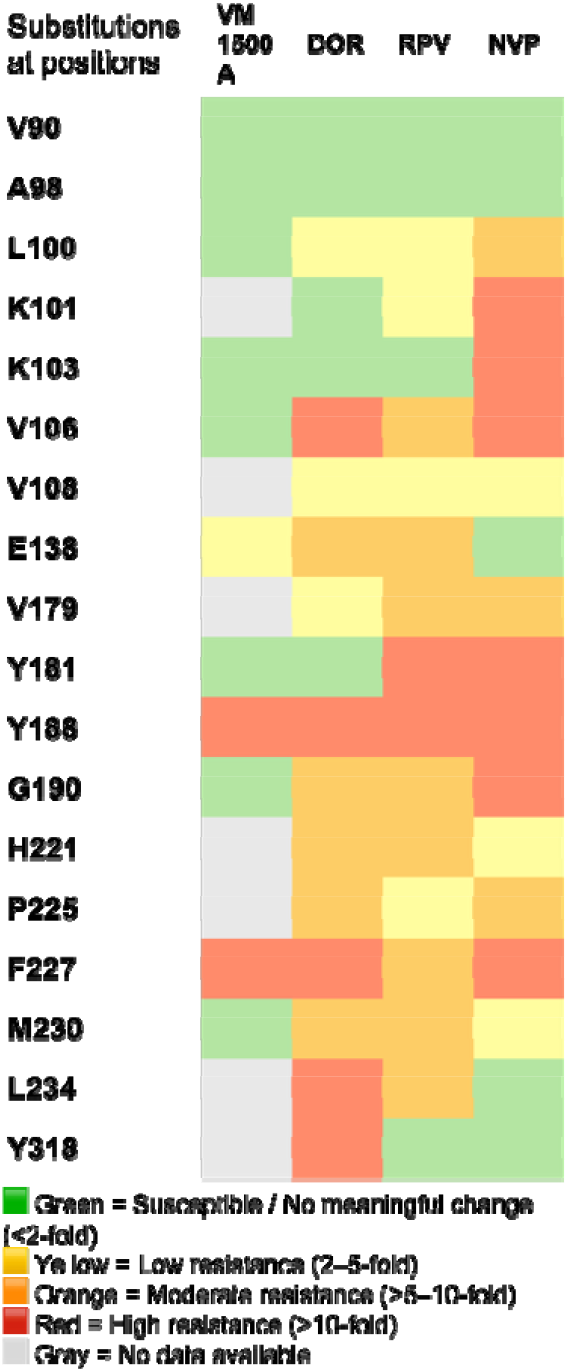
NNRTI resistance profile comparison across common HIV-1 reverse transcriptase mutations. Heatmap summarizing the resistance profiles of VM1500A/depulfavirine, doravirine (DOR), rilpivirine (RPV), and nevirapine (NVP) against a panel of clinically relevant HIV-1 reverse transcriptase mutations. Resistance levels are categorized based on the fold change in EC_50_ values relative to the wild-type virus. Green indicates susceptible or no meaningful change (<2-fold), yellow indicates low-level resistance (2–5-fold), orange indicates moderate resistance (>5–10-fold), and red indicates high-level resistance (>10-fold).

Similarly, DOR also exhibits high-level resistance to mutations such as Y188L, F227C, and the V106A/F227C combination. Additional DOR resistance-associated mutations, including L234I and Y318F, have not yet been evaluated with elsulfavirine (ESV) or depulfavirine. It remains unclear whether these mutations will emerge during ESV treatment, as this antiviral is relatively new and clinical data remain limited. Moderate reductions in DOR susceptibility have been reported for mutations including L100I, V106A, V108I, E138K, G190E/S, H221Y, and M230I/L (24, 32-36). RPV demonstrates variable reductions in susceptibility across several common NNRTI resistance mutations, including K101E, E138K, and Y181C, as well as Y188L and F227C (35, 37). Additional moderate resistance to RPV has been observed with V106A, V179D/I/F, G190E, H221Y, and M230I/L (35, 37). NVP exhibits the most limited resistance profile among the inhibitors compared, consistent with the rigidity of its molecular scaffold. It displays high-level resistance with mutations such as K103N, V106A, Y181C, Y188L, and G190A, with additional moderate resistance associated with V179D/I/F, P225H, and M230I/L (Figure 4) (35, 38).

The resistance pattern observed for depulfavirine can be explained by its distinct structural footprint and the specific interactions it forms within the NNRTI-binding pocket. Depulfavirine adopts an extended three-ring conformation that reaches deeper into the hydrophobic tunnel than many other NNRTIs, forming additional stabilizing contacts that support activity against common resistance mutations. Specifically, depulfavirine forms a strong π–π interaction with Y188 and engages residues near the pocket entrance, including K101–S105. These interactions provide alternative anchoring points that help buffer local structural perturbations caused by single-amino-acid substitutions. Consistent with this binding mode, depulfavirine retains activity against mutations at positions 101, 103, and 190 because it engages both the entrance and deeper regions of the NNRTI-binding pocket. Although K103 lies at the pocket entrance, depulfavirine interacts primarily with the backbone rather than the side chain, which may help preserve activity against K103N and other mutations. Similar to K103N, Y181C has minimal impact on depulfavirine potency because Y181 adopts an alternate conformation that points away from the inhibitor, resulting in limited direct contact.

High-level resistance to depulfavirine emerges primarily with Y188L, F227C (Supplemental Figure 1), or V106A/F227C (11). Y188L likely disrupts the π–π stacking interaction that stabilizes depulfavirine binding. In contrast, the combination of F227C with V106A may alter the hydrophobic environment or geometry of the tunnel, preventing optimal positioning of the extended scaffold. Notably, these mutations that reduce depulfavirine potency also impair viral fitness, as reported previously (39, 40). The requirement for multiple coordinated changes reflects depulfavirine’s broad interaction footprint and supports its higher genetic barrier to resistance than that of other NNRTIs. *In vitro* RT primer extension assays were performed to evaluate the inhibitory potency of depulfavirine against wild-type HIV-1 RT and a panel of clinically relevant NNRTI-resistant mutant enzymes (Figure 5, Sup Fig 5). Depulfavirine demonstrated activity against wild-type RT, with an IC_50_ in the high-nanomolar range, slightly higher than previously reported antiviral potency (Figure 5) (11). This could be due to several factors, including the experimental setup (limited portion of the full HIV-1 replication cycle) and multiple target effects. Across most single resistance substitutions, including K103N, V106A, Y181C, F227C, and E138K. Depulfavirine retained activity with the greatest increase in IC_50_ values associated with the F227C single and double mutant F227C V106A. These results indicate that many common NNRTI resistance mutations have a limited impact on depulfavirine binding or enzymatic inhibition. Of note, Y188L was not tested. This pattern aligns with the resistance profile observed in cell-based assays and highlights a narrow set of structural vulnerabilities within the NNRTI-binding pocket conferring resistance to similar compounds. Collectively, the IC_50_ results support the biochemical robustness of depulfavirine and its capacity to maintain inhibitory activity across a wide range of NNRTI-resistant variants.

**Figure 5.**
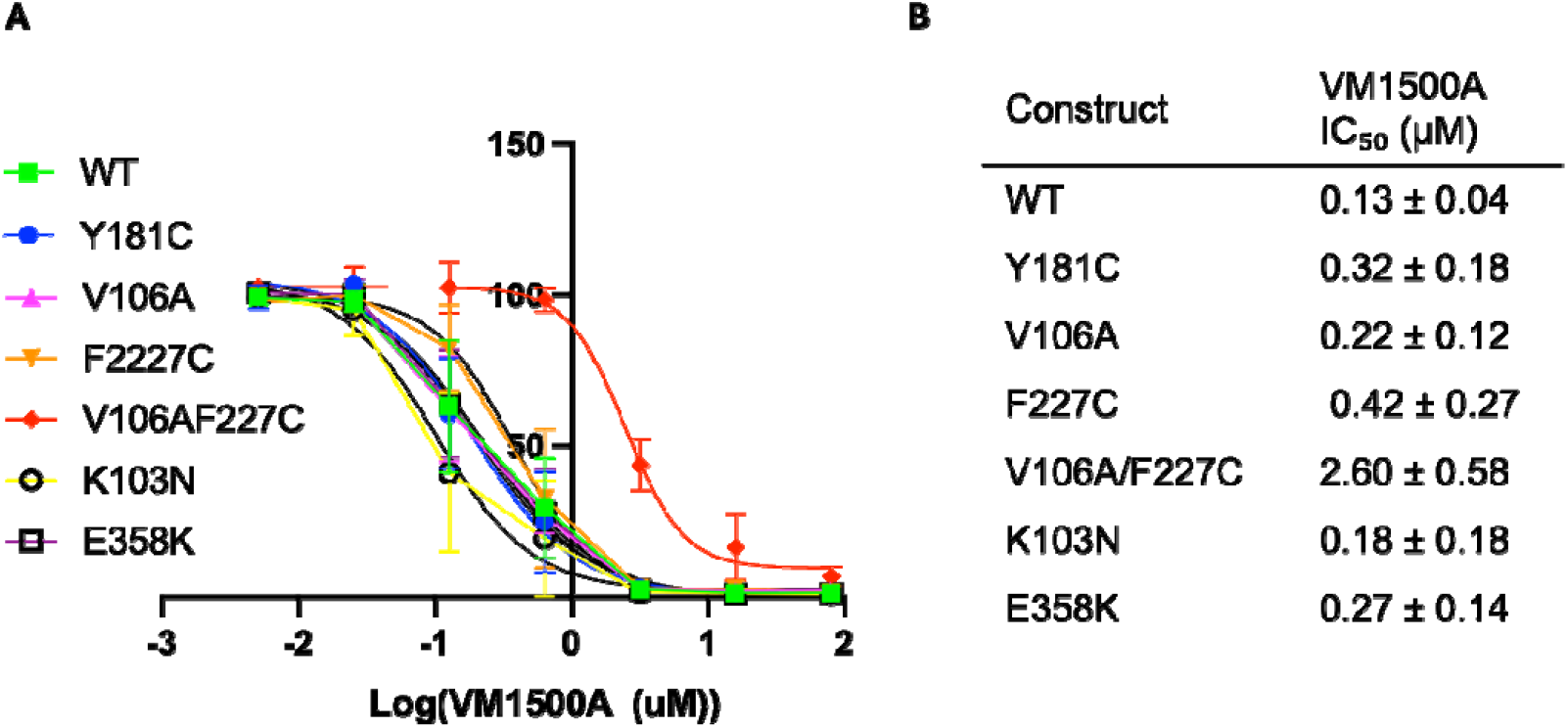
Inhibition of NNRTI-resistant HIV-1 RT enzymes by depulfavirine in primer extension assays. Primer extension reactions were performed using WT and NNRTI-resistant RT enzymes with the Td31/Pd18-P0 substrate. Reactions were carried out for 15 minutes in the presence of 1 μM dNTPs, MgCl_2_, and increasing concentrations of depulfavirine (0–400 μM). Extension activity was quantified and normalized to no-drug controls. Data represent mean ± SD from independent experiments.

### Synergy

Given depulfavirine’s favorable resistance profile, biochemical properties, and long-acting potential, we evaluated its antiviral synergy with other long-acting antiretroviral agents and/or standards of care (Figure 6, Sup Figure 2 and 3). To evaluate potential long-acting combination partners for elsulfavirine (ESV), antiviral activity was measured across dose-response matrices in cell-based assays using islatravir (ISL), cabotegravir (CAB), lenacapavir (LEN), and tenofovir disoproxil fumarate (TDF). ESV, the prodrug of depulfavirine, was selected for these experiments based on its favorable solubility and cell permeability. The partner antiretrovirals represent diverse mechanisms of action and include agents with established or emerging long-acting utility: islatravir, a nucleoside reverse transcriptase translocation inhibitor (NRTTI); cabotegravir, a long-acting integrase strand transfer inhibitor; lenacapavir, a long-acting capsid inhibitor; and TDF, a widely used nucleoside reverse transcriptase inhibitor currently being explored in long-acting formulations.

**Figure 6.**
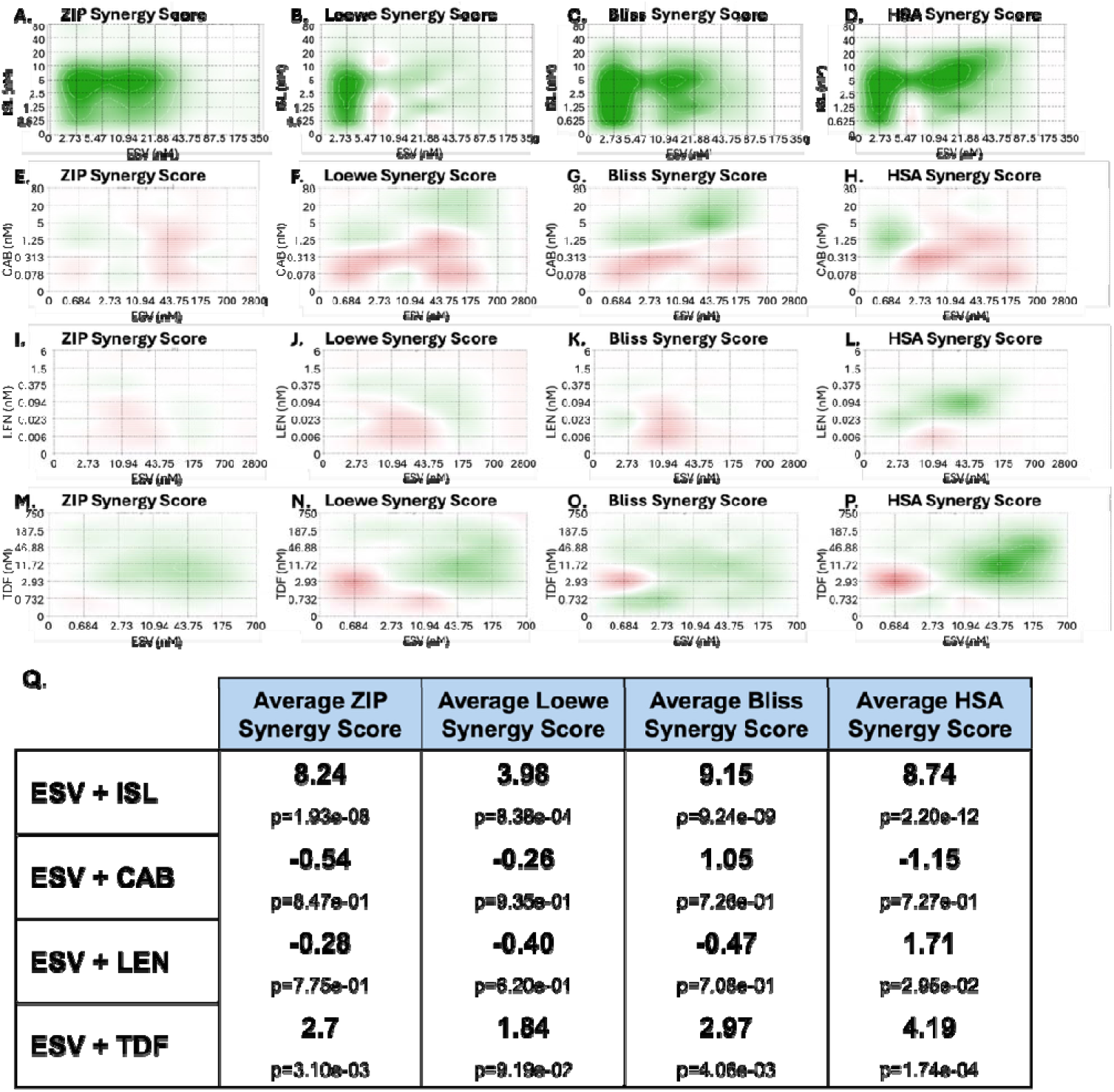
Synergy analysis of ESV in combination with long-acting antiviral candidates. Synergy scores were calculated at each dose–response point and as overall synergy scores using the ZIP, Loewe, Bliss, and HSA models. Drug combinations tested included ESV+ISL, ESV+CAB, ESV+LEN, and ESV+TDF. Percent inhibition values were calculated and analyzed using SynergyFinder Plus. Two-dimensional synergy plots are shown in panels A–P, with synergy scores summarized in table format in panel Q (18).

Drug interactions were quantified using four established statistical models implemented in the SynergyFinder+ platform: Highest Single Agent (HSA), Bliss Independence (Bliss), Loewe Additivity (Loewe), and Zero Interaction Potency (ZIP). Each model assumes a null hypothesis of non-interaction, where scores above zero indicate synergy and scores below zero indicate antagonism; because these models differ in their underlying assumptions, they may yield divergent interpretations of the same dataset (23). For this analysis, synergy was defined as a score greater than 10, antagonism as a score below -10, and scores between -10 and 10 were considered additive.

The HSA model, frequently applied in cancer drug research, defines synergy as a combination effect exceeding the maximum effect of either agent alone, making it particularly useful when no single drug achieves complete inhibition (41). The Bliss Independence model assumes that each drug acts through independent pathways, such that the expected combined response is the multiplicative product of the individual response probabilities (42). The Loewe Additivity model treats the two drugs as functionally equivalent, defining the additive baseline as the expected response of each drug combined with itself; deviations from this baseline indicate synergy or antagonism (43). The ZIP model integrates the principles of both Bliss and Loewe by assuming that drugs act independently and that neither drug shifts the potency of the other. A delta score is derived from the deviation of the observed combination response from the expected individual dose-response curves (44). Synergy landscapes were visualized across the full concentration matrices (Figure 6). SynergyFinder+ calculates synergy scores at each dose-response point, which are plotted into synergy matrices for each model along with mean synergy scores and corresponding p-values for each combination (Zheng et al., 2022).

Among the combinations tested, ESV + ISL demonstrated the strongest interaction, showing near-synergistic additivity (synergy defined as >10). Positive synergy scores were observed across all four models, with mean ZIP, Loewe, Bliss, and HSA scores of 8.24, 3.98, 9.15, and 8.74, respectively. These effects were statistically significant across models (p ≤ 10^−3^ to 10^−12^), and the synergy landscape revealed broad additivity and synergy across the dose matrix, indicating additive antiviral activity across multiple concentration combinations. This may be due to the formation of a dead-end complex, as previously described for other NRTI+NNRTI combinations (45). This pairing may be of particular interest given that ISL may have a complementary resistance profile to depulfavirine with viruses containing F227 substitutions (36, 46). ESV + CAB showed largely additive behavior, with mean synergy scores near zero across all models (ZIP: -0.54, Loewe: -0.26, Bliss: 1.05, HSA: -1.15). Similarly, ESV + LEN exhibited predominantly additive interactions, with mean synergy scores close to zero across the ZIP, Loewe, Bliss, and HSA models (-0.28, -0.40, -0.47, and 1.71, respectively). ESV + TDF displayed moderate additive activity, with positive mean scores across all four models (ZIP 2.7, Loewe 1.84, Bliss 2.97, HSA 4.19). These findings support the compatibility of ESV with multiple classes of antiretroviral agents and highlight ISL as a potential partner for long-acting combination strategies.

## Conclusion

Our structural, biochemical, and synergy analyses demonstrate that depulfavirine adopts a unique binding mode within the NNRTI-binding pocket that underlies its broad activity against a wide range of clinically relevant NNRTI resistance mutations. The extended three-ring scaffold, deep hydrophobic engagement, and additional interactions at both the entrance and base of the pocket collectively confer a higher genetic barrier to resistance than some currently approved NNRTIs. Consistent with these structural features, depulfavirine maintains nanomolar IC_50_ values across most single-resistance substitutions, with substantial losses in potency limited to F227C and the combined V106A/F227C pathway, mutations that also reduce viral fitness. In cell-based assays, ESV demonstrated favorable pairing with multiple long-acting antiretroviral agents, supporting its potential compatibility within future long-acting combination regimens. Together, these findings highlight depulfavirine as a next-generation NNRTI with a favorable resistance profile, combination potential, and long-acting profile.

