## Supplemental Figure 1 for "HIV-1 Reverse Transcriptase interactions with Long-acting NNRTI, Depulfavirine (VM1500A)"

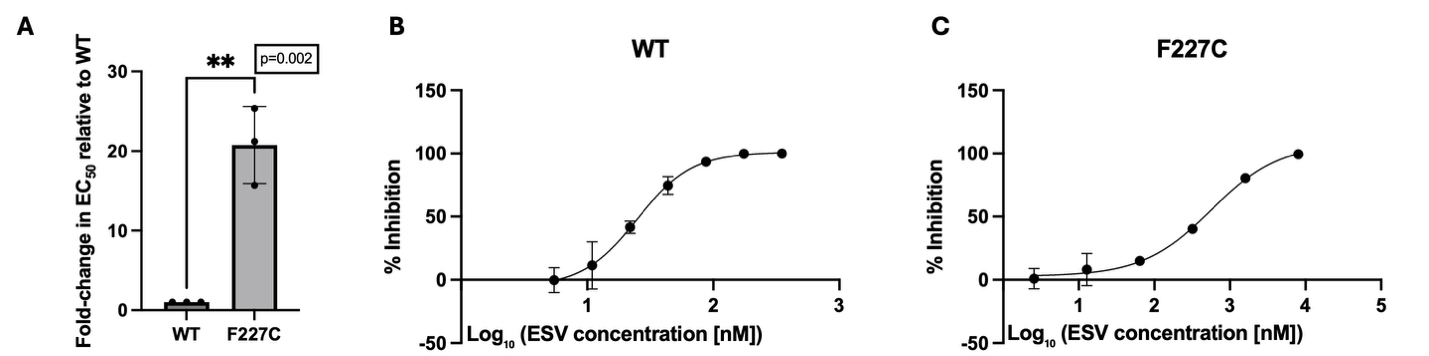

**Supplemental Figure 1.** Dose–response curve of ESV viral inhibition in WT and F227C HIV-1. Dose–response curves showing inhibition of HIV-1 infection by ESV across increasing concentrations for WT and F227C variants. TZM-GFP cells were pre-incubated with ESV for 24 h before infection with NL4-3 Δenv VSVg pseudotyped virus. Infection levels were quantified by GFP expression 48 h post-infection. Percent inhibition was calculated relative to untreated control wells and plotted against log₁₀-transformed ESV concentrations (nM). Data were fit using a four-parameter nonlinear regression model in [GraphPad Prism](https://www.graphpad.com/?utm_source=chatgpt.com) to determine EC₅₀ values. Relative fold-change in EC₅₀ values (WT vs F227C) was calculated. Error bars represent mean ± SEM. Statistical significance between WT and F227C EC₅₀ values was determined using an unpaired two-tailed t-test.
