## Supplemental Figure 2 for "HIV-1 Reverse Transcriptase interactions with Long-acting NNRTI, Depulfavirine (VM1500A)"

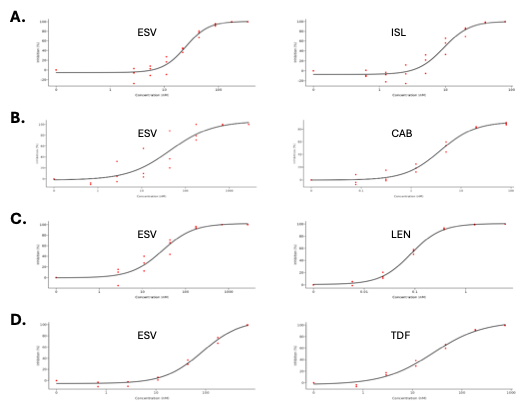

**Supplemental Figure 2.** Dose–response and inhibition matrices from combination treatments against HIV-1. Dose–response curves for ESV in combination with (A) ISL, (B) CAB, (C) LEN, and (D) TDF. Assays were performed in TZM-GFP cells, with dose–response matrices pre-incubated for 24 h before infection with NL4-3 Δenv pseudotyped virus. GFP expression was imaged 48 h post-infection, and percent inhibition values were calculated and analyzed using SynergyFinder Plus.
