## Supplemental Figure 3 for "HIV-1 Reverse Transcriptase interactions with Long-acting NNRTI, Depulfavirine (VM1500A)"

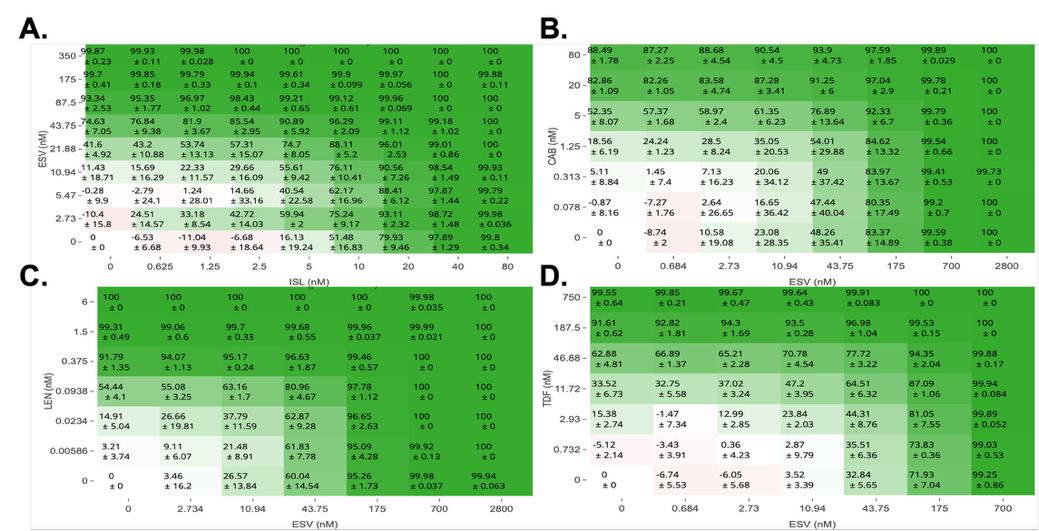

**Supplemental Figure 3.** Representative percent inhibition matrices for ESV combination treatments. The matrices show percent inhibition for ESV in combination with (A) ISL, (B) CAB, (C) LEN, and (D) TDF, calculated using SynergyFinder Plus.
