## Supplemental Figure 4 for "HIV-1 Reverse Transcriptase interactions with Long-acting NNRTI, Depulfavirine (VM1500A)"

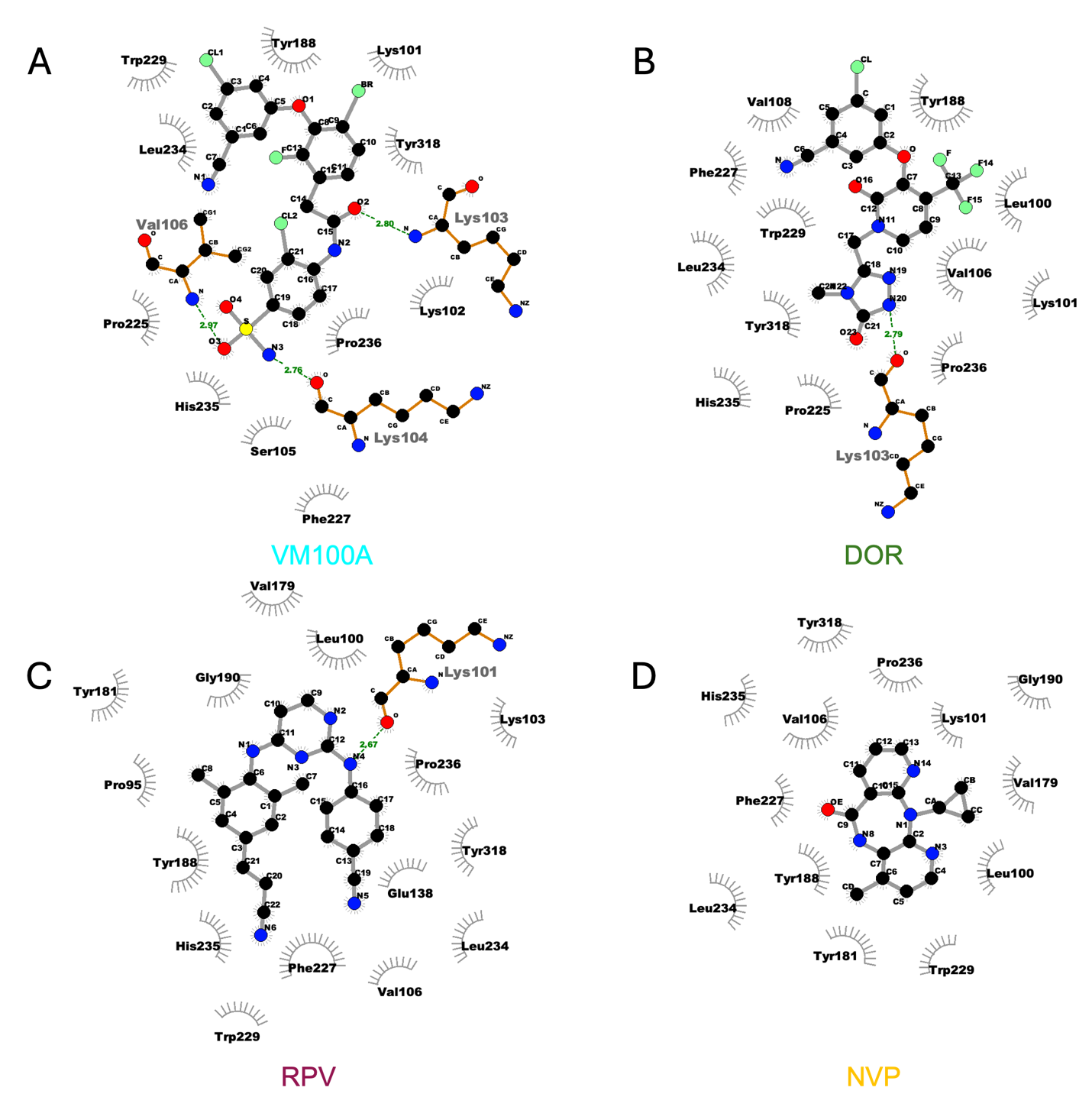

**Supplemental Figure 4.** Ligand–protein interaction diagrams of NNRTIs bound to HIV-1 reverse transcriptase. Two-dimensional interaction plots of (A) depulfavirine (VM1500A), (B) doravirine (DOR), (C) rilpivirine (RPV), and (D) nevirapine (NVP) in complex with HIV-1 reverse transcriptase, generated using LigPlot+. Hydrogen bonds are shown as dashed lines, and hydrophobic interactions are represented by spoked arcs. Key interacting residues within the NNRTI binding pocket are labeled. These diagrams highlight differences in binding interactions and contact profiles among NNRTIs. LigPlot+ diagrams were generated using LigPlot+ software (47).
