## Supplemental Figure 5 for "HIV-1 Reverse Transcriptase interactions with Long-acting NNRTI, Depulfavirine (VM1500A)"

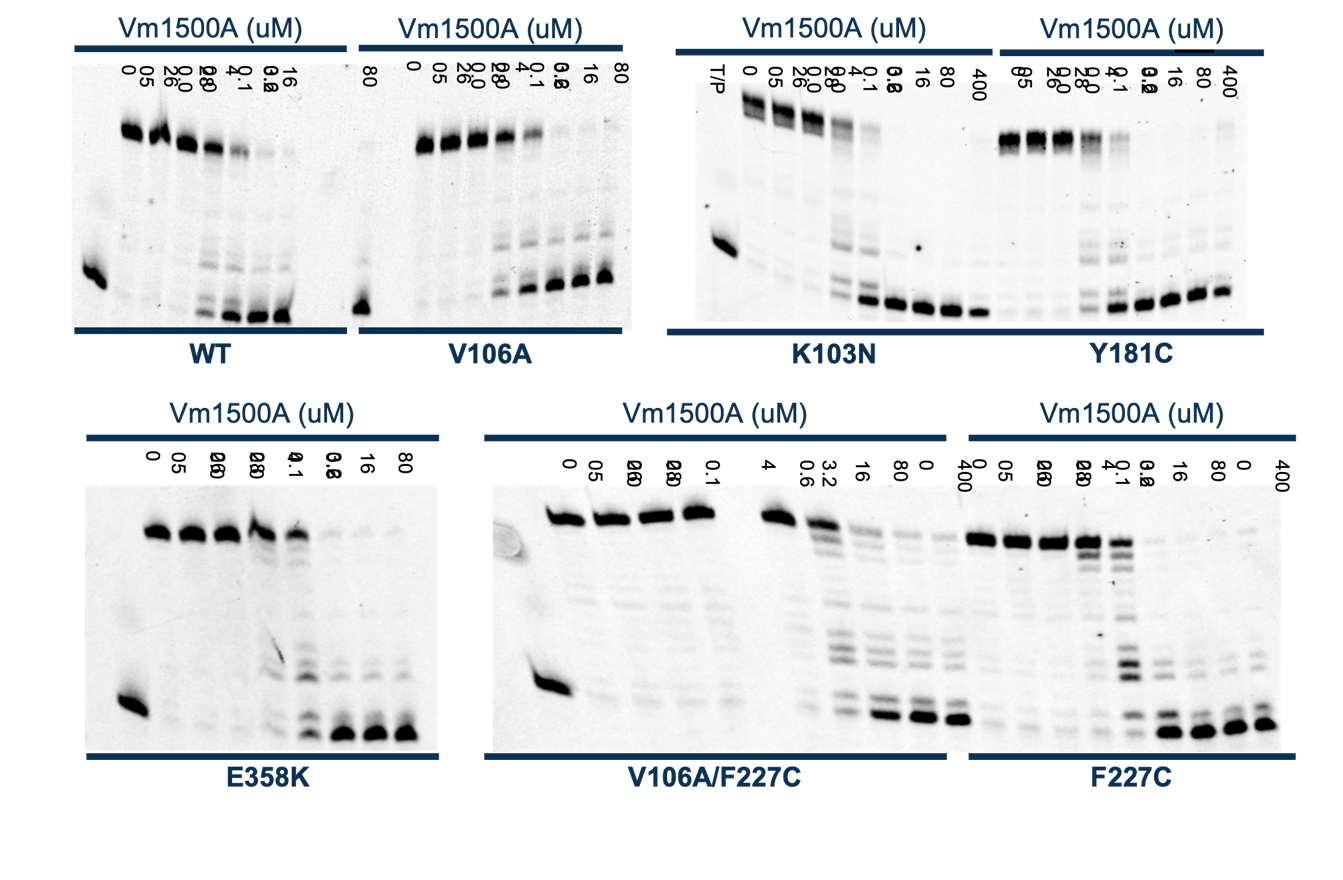

**Supplemental Figure 5**. Primer extension assays of NNRTI-resistant RT enzymes inhibited by depulfavirine. T_d31_/P_d18-P0_ was incubated with WT or mutant RT for 15 minutes in the presence of 1 μM dNTPs, MgCl_2_, and increasing concentrations of depulfavirine (0-400 µM).
