## Supplemental Table 1 for "HIV-1 Reverse Transcriptase interactions with Long-acting NNRTI, Depulfavirine (VM1500A)"

**Supplemental Table 1. Data collection and refinement statistics (PDB: 7TAZ).**

| **Parameter** | **7TAZ** |
| --- | --- |
| **Wavelength** | 1.0 |
| **Resolution range (Å)** | 31.38–2.40 (2.49–2.40) |
| **Space group** | C 1 2 1 |
| **Unit cell (Å, °)** | 163.05, 73.87, 108.64; 90, 99.96, 90 |
| **Total reflections** | 318,061 |
| **Unique reflections** | 46,997 (3,443) |
| **Multiplicity** | 6.8 (6.9) |
| **Completeness (%)** | 99.9 (100.0) |
| **Mean I/σ(I)** | 20.32 (3.73) |
| **Wilson B-factor** | 65.78 |
| **R-merge (%)** | 4.7 (68.9) |
| **R-meas (%)** | 5.2 (74.4) |
| **R-pim (%)** | 2.0 (28.4) |
| **CC1/2 (%)** | 99.9 (90.6) |
| **Reflections used in refinement** | 49,930 (4,972) |
| **Reflections used for R-free** | 2,472 (242) |
| **R-work** | 0.2487 (0.3269) |
| **R-free** | 0.2774 (0.3586) |
| **Number of non-hydrogen atoms** | 7,804 |
| macromolecules | 7,687 |
| ligands | 46 |
| solvent | 84 |
| **Protein residues** | 934 |
| **RMS(bonds)** | 0.004 Å |
| **RMS(angles)** | 0.80° |
| **Ramachandran favored (%)** | 94.46 |
| **Ramachandran allowed (%)** | 4.89 |
| **Ramachandran outliers (%)** | 0.65 |
| **Rotamer outliers (%)** | 2.61 |
| **Clashscore** | 8.45 |
| **Average B-factor** | 90.76 |
| macromolecules | 91.05 |
| ligands | 86.38 |
| solvent | 66.27 |
